# Comprehensive phylogenetic reconstructions support ancestral omnivory in the ecologically diverse bat family Phyllostomidae

**DOI:** 10.1101/2025.02.04.636560

**Authors:** Xueling Yi, Dimitrios-Georgios Kontopoulos, Michael Hiller

## Abstract

Adaptive radiations often occur with an early burst of ecological diversification, which requires not only various available niches but also a generalist ancestor with wide ecological niche breadths. However, ancestral generalism remains hard to test in empirical cases. The New World leaf-nosed bats (family Phyllostomidae) represent an unparalleled mammalian adaptive radiation with diverse dietary niches including arthropods, blood, terrestrial vertebrates, nectar, and fruits. However, when and how often phyllostomid bats transitioned from insectivory to fruit or nectar feeding remains unclear. Here we tested the hypotheses of ancestral insectivory versus ancestral omnivory in Phyllostomidae (141 species) using improved trait reconstructions based on multi-response phylogenetic threshold models, while explicitly accounting for phylogenetic uncertainty. Our results indicate that complementary fruit feeding has fully evolved at the early burst of the phyllostomid radiation and started to evolve in the most recent common ancestor of the family, supporting the ancestral omnivory hypothesis. In addition, fruit feeding probably evolved before nectar eating in Phyllostomidae, in contrast to the claims of previous studies. Extending this analysis to all bat families (621 species) reveals independent evolution of ancestral fruit feeding in four families, namely the Pteropodidae (Old World fruit bats) and three families from the Noctilionoidea superfamily. Despite the ancestral omnivory of these fruit-eating families, only Phyllostomidae and Pteropodidae show high species diversity and evolved predominant and strict fruit feeding. Therefore, our results reveal that ancestral generalism (i.e., omnivory) may be a precondition of but does not necessarily lead to adaptive radiations which also require subsequent niche partitioning and speciation.

## Introduction

Adaptive radiation refers to the evolutionary process of an ancestor generating diverse descendant species adapted to various ecological niches. The major invasion of diverse niches is proposed to occur soon after the beginning of the radiation, leading to an “early burst” of speciation and ecological variation (Gavrilets & Losos, 2009). This early burst stage likely not only requires access to diverse niches (e.g., ecological opportunities) but also generalism of the common ancestor that have expanded niche breadths (Gillespie et al., 2020; Losos, 2010; Schluter, 2000). The early burst pattern of adaptive radiation has been supported by several empirical studies using eco-morphological traits, including body shape in the Tanganyikan cichlid fishes (Ronco et al., 2021), bill diversity in birds (Cooney et al., 2017), and diet-related dental traits in Carnivora (Slater & Friscia, 2019) and the New World leaf-nosed bats (Grossnickle et al., 2024). However, the generalism of the ancestor remains often unclear and difficult to test, especially in species lacking fossil data, such as bats (Brown et al., 2019).

A great study system to investigate adaptive radiations and ancestral generalism are bats, the second largest mammalian order (Chiroptera) with >1480 recognized species (Simmons & Cirranello, 2024) distributed globally in various habitats. The New World leaf-nosed bats (family Phyllostomidae) are the most ecologically diverse bat family that originated from an adaptive radiation. While most bats are strict insectivores, phyllostomid bats have radiated into 11 subfamilies with 227 recognized species that feed on various diets including arthropods, blood, terrestrial vertebrates, fruits, and nectar. This extremely high dietary diversity is unparalleled in mammals. Key to the phyllostomid adaptive radiation is the ability to feed on fruits and nectar which are found as complementary or major food sources in many species (Rex et al., 2010; Rojas et al., 2018). Specialized fruit and nectar feeding has been associated with main morphological diversification (Dumont et al., 2012; Freeman, 2000) and higher speciation rates (Rojas et al., 2018; Shi & Rabosky, 2015).

Consistent with the hypothesis of ancestral generalism, previous studies have proposed that the ancestors of Phyllostomidae were omnivorous that already included fruits in addition to insects in their diet (Baker et al., 2012; Freeman, 2000), which might have facilitated their initial invasion of alternative dietary niches. This ancestral omnivory hypothesis (i.e., ancestral generalism; note that “omnivory” here only refers to diet diversity rather than equal proportions of plant- and animal-based food) has been supported by the inferred diversity of ancestral sensory structures (Hall et al., 2021; Mutumi et al., 2023), patterns of selection of sensory and metabolic genes on the ancestral phyllostomid branch (Davies et al., 2020; Potter et al., 2021), and the early burst pattern found using diet-related molar traits (Grossnickle et al., 2024). In contrast, previous diet reconstructions supported the alternative ancestral insectivory hypothesis and inferred multiple independent transitions from insect feeding to fruit or nectar feeding during the phyllostomid radiation (Rojas et al., 2011). Another reconstruction showed low probabilities of nectar feeding at the early burst stage, but still indicated insect feeding as the most likely ancestral state and relatively later evolution of fruit feeding in Phyllostomidae (Grossnickle et al., 2024).

Previous diet reconstructions have several limitations. First, previous reconstructions merged all diets into one single trait and coded each species only by their predominant diet while ignoring complementary food items (Grossnickle et al., 2024; Rojas et al., 2011). This practice necessarily eliminates part of the complexity in diet evolution, which can affect ancestral reconstructions, potentially masking the signal of omnivory. Indeed, when nectar and fruit feeding were modelled as independent traits, the reconstructions indicated ancestral omnivory although the results had high uncertainty (Rojas et al., 2011). Second, the constraints enforced in evolutionary models used in previous reconstructions may have biased the results. For example, the ordered model in (Rojas et al., 2011) specified transitions from insectivory to other diets, thus enforcing ancestral insectivory. The model in (Grossnickle et al., 2024) used equal transition rates among arthropod, fruit or nectar diets, which may be an overly simplifying assumption at best. Third, phylogenetic uncertainty needs to be considered because several nodes in the phyllostomid phylogeny remain unresolved (Camacho et al., 2022; Grossnickle et al., 2024; Rojas et al., 2011, 2016; Shi & Rabosky, 2015), especially regarding the positions of the nectar-feeding subfamily Glossophaginae, the omnivorous subfamily Phyllostominae, and the insect-feeding subfamily Lonchorhininae (Fig. 1A). Because the most recent common ancestor (MRCA) of these subfamilies represents the early burst stage (Grossnickle et al., 2024) when major diet diversification and rapid speciation occurred, diet reconstructions relying on a single phylogenetic tree may not be accurate, and phylogenetic uncertainty needs to be taken into account.

**Figure 1.**
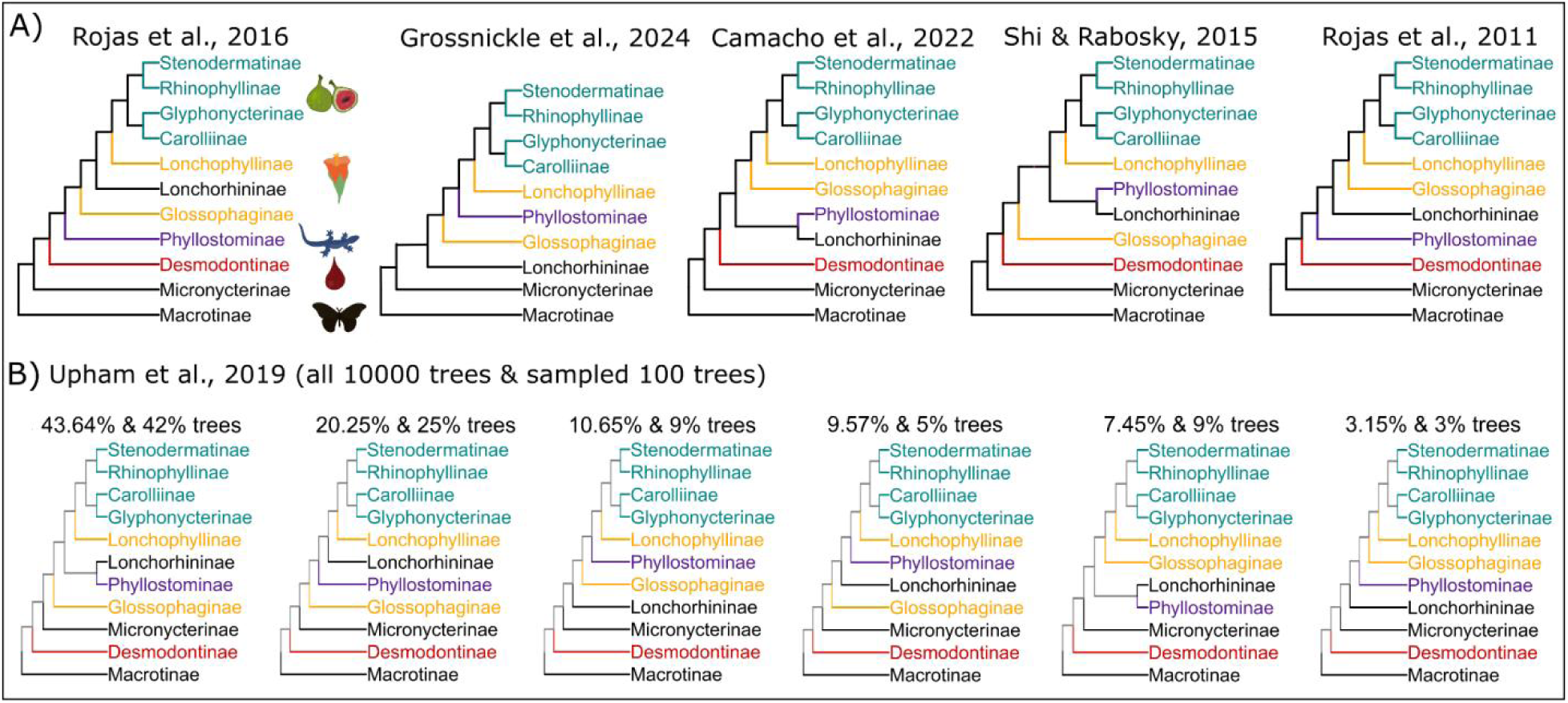
Major subfamily topologies of Phyllostomidae in A) the literature and B) the analyzed posterior distribution of phylogenetic trees. Subfamilies are colored by their predominant diets. **A)** The topology in (Rojas et al., 2016) has been considered the most reliable and is supported by phylogenomic analyses. The vampire bats (subfamily Desmodontinae) were excluded in (Grossnickle et al., 2024). **B)** (Upham et al., 2019) inferred 10,000 mammalian phylogenetic trees using an assembled 31-gene supermatrix and Bayesian inference. The panel shows the six most prevalent phyllostomid subfamily topologies that together cover 94.71% of the 10,000 trees and 93% of the 100 trees sampled in this study. Corresponding percentages of each topology are shown on top of the tree.

Here, we tested the hypotheses of ancestral generalism (i.e., omnivory) versus insectivory in adaptive radiations using phyllostomid bats as the study system. The ancestral omnivory hypothesis proposes the evolution of complementary fruit/nectar feeding no later than the early burst stage, whereas the ancestral insectivory hypothesis proposes multiple more recent transitions from insect feeding to fruit and nectar feeding. In addition, in contrast to previous reconstructions (Grossnickle et al., 2024), we hypothesize that fruit feeding evolved earlier than the presumably more specialized nectar feeding. To overcome the above-mentioned limitations in previous studies, first, we treated each diet as a distinct trait, coded as four ordered discrete states representing its relative importance in the species diet (absent, complementary, predominant, strict). We combined all traits into one generalized linear mixed model fitted by Bayesian Markov Chain Monte Carlo (MCMC) implemented in the R package MCMCglmm (Hadfield, 2015). Second, we fitted trait evolution using the threshold model that assumes an unknown continuous variable (liability) underlying the ordered states (Felsenstein, 2012). Under the threshold model, traits can only change between adjacent states, and reverse state changes are more likely to happen sooner than later after the first state change (Revell, 2014), which makes the threshold model more biologically meaningful than the traditional Markov chain model used in previous studies (Grossnickle et al., 2024; Rojas et al., 2011). Third, to account for phylogenetic uncertainty, we incorporated a posterior distribution of phylogenies and also considered the most likely subfamily topology (Rojas et al., 2016) which is supported by phylogenomics. We found early evolution of complementary fruit feeding inferred both in the early burst stage and the common ancestor of Phyllostomidae (analyses of 176 species), supporting the hypothesis of ancestral omnivory and thus generalism. Extending this analysis to all bat families (621 species) indicated additional independent evolution of ancestral fruit feeding in another four families including the Old World fruit bats (Pteropodidae).

## Materials and Methods

### Taxonomy and phylogenetic trees

To cover the phyllostomid diversity and account for phylogenetic uncertainty, we used a posterior distribution of mammalian phylogenetic trees generated in a Bayesian framework (MrBayes; (Upham et al., 2019)). To this end, we downloaded from VertLife (http://vertlife.org/phylosubsets) all 10,000 node-dated trees inferred using only taxa with DNA sequences (i.e., the “DNA-only” trees), following the recommendation of (Upham et al., 2019) for trait reconstructions. We additionally excluded the species having only one gene sequence because the lack of data in these species likely added more noise than real phylogenetic uncertainty. This retained a total of 624 bats covering all 21 bat families, all 11 phyllostomid subfamilies, and 55 of the 61 phyllostomid genera (species taxonomy updated based on https://batnames.org/, accessed on 2024-10-16; (Simmons & Cirranello, 2024)). For the analyses of Phyllostomidae, we pruned these trees by keeping all species in the family Phyllostomidae (n=141) and the other five closely related families in the superfamily Noctilionoidea (n=8 in Mormoopidae, n=2 in Noctilionidae, n=1 in Furipteridae, n=2 in Thyropteridae, and n=1 in Mystacinidae). In addition, we kept several outgroup species fromthree more insectivorous families (n=17 in Vespertilionidae, n=3 in Molossidae, n=4 in Emballonuridae) in the suborder Yangochiroptera. Fish-feeding *Myotis* bats were particularly selected to be part of the outgroup to maximize the representation of this diet.

Since these phylogenetic trees were inferred from only up to 31 genes (Upham et al., 2019), we examined how reliable these trees are by checking the monophyly of each bat family and phyllostomid subfamily across all 10,000 trees. We also tested the monophyly of several internal nodes, such as the phyllostomid early burst node representing the MRCA of 8 subfamilies excluding Desmodontinae, Micronycterinae, and Macrotinae. The early burst node in (Grossnickle et al., 2024) was placed after the divergence of the subfamily Lonchorhininae, but because the position of this subfamily remained unresolved (Fig. 1A), we labeled the early burst node before its divergence. After confirming a high degree of monophyly of bat families and phyllostomid subfamilies in these 10,000 trees, we estimated frequencies of subfamily-level topologies in all 10,000 trees and in 100 trees that were randomly sampled as a representative dataset of the posterior distribution. We also generated a dataset by selecting all trees that show the subfamily topology found in (Rojas et al., 2016), which is consistent with phylogenomic analyses of high-quality genomes and thus likely represents the true subfamily relationships.

### Bat diets

We used the diet database of the superfamily Noctilionoidea (Rojas et al., 2018) and incorporated diets of other bats from EltonTraits 1.0 (Wilman et al., 2014) and from publications ((Castillo-Figueroa et al., 2022; Frick et al., 2009, 2014; López-Aguirre et al., 2022; Santana et al., 2011); see notes in Table S1). A total of eight diets were recorded in (Rojas et al., 2018), each coded in four ordered states representing the relative importance in the overall species diet: 0 for absent component, 1 for complementary (defined as less than 40% in the overall diet), 2 for predominant (more than 60%), and 3 for strict (the only component in the diet). For three of the eight diets, only two states were observed among bats (binary diets). Among them, blood feeding only exists in vampire bats. Seed feeding was only recorded in two species (*Chiroderma doriae*, *Chiroderma villosum*) that are also predominantly fruit-feeding. Similarly, feeding on leaves and flower pieces was only recorded as complementary in species that also feed on nectar and/or fruits. Because these binary traits are only restricted to few groups, we excluded these three binary diets, leaving five diets (arthropods, terrestrial vertebrates, fish, pollen and nectar, fruits) in the main analyses.

The above-incorporated diet data (hereafter the raw coding) is overall inclusive. For example, species in the basal phyllostomid subfamilies (Macrotinae and Micronycterinae) were coded as complementary fruit feeding (Rojas et al., 2018) although the majority of literature considers them as insectivorous (e.g., (Fleming et al., 2020; Grossnickle et al., 2024)). Similarly, the subfamily Lonchorhininae has been considered strictly insectivorous whereas a species, *Lonchorhina aurita*, was coded as complementary fruit-feeding (Rojas et al., 2018). We reviewed the literature on these three subfamilies and found robust support for complementary fruit feeding in a few species of the subfamily Micronycterinae (*Lampronycteris brachyotis*, *Micronycteris hirsuta,* and *Micronycteris minuta*; (Bonaccorso, 1979; Gardner, 1977; Giannini & Kalko, 2004; Humphrey et al., 1983)). However, we found little dietary information and/or strongly supported insect feeding but very sparse and ambiguous descriptions of fruit feeding in the other eight species, including five *Micronycteris* species (*M. megalotis*, *M. nicefori*, *M. schmidtorum*, *M. sylvestris*, *M. microtis* (Gardner, 1977; Giannini & Kalko, 2004; Howell & Burch, 1973; Humphrey et al., 1983, p. 198; Kalka & Kalko, 2006; Rojas et al., 2011; Santana et al., 2011)), two *Macrotus* species (*M. waterhousii*, *M. californicus* (Anderson, 1969; Gardner, 1977; Sánchez et al., 2016)), and *Lonchorhina aurita* ((Lassieur & Wilson, 1989); details of the diet review is in Table S2). To explore the robustness of our diet reconstructions at phyllostomid basal nodes, we also fit the models using manually re-coded input diets (hereafter the basal-insect coding), where these eight species were changed to absent fruit-feeding and strict arthropod-feeding (except for *Micronycteris microtis*, which was kept as predominant arthropod-feeding because of the complementary vertebrate-feeding (Santana et al., 2011)).

### Ancestral reconstruction of ordered diet states under the threshold model

The five diets were included as response variables in a Markov chain Monte Carlo multi-response generalized linear mixed model fitted using the R package MCMCglmm (Hadfield, 2015). All analyses in this study were done in the R version 4.4 (R Core Team, 2024). The scripts to fit MCMCglmm models were modified from (Kontopoulos et al., 2025). The model includes one distinct intercept for each response variable and integrates the phylogeny as a random effect on the intercepts using the inverse of the phylogenetic covariance matrix extracted with the *inverseA* function. Because the residual variance is not identifiable in threshold models, we fixed it to 1. We also forced the threshold liabilities to range between -7 and 7 to facilitate their estimation (trunc=TRUE). To minimize the influence of the prior distributions, we used relatively uninformative parameter expanded priors (nu=5; nu=6 when including vampire bats). For each phylogenetic tree, MCMCglmm was run with 10 independent chains, each chain with a total of two million iterations, 10% (200000) burnin, and a sampling interval of 50 iterations. Convergence of all 10 chains for each phylogenetic tree was confirmed based on the effective sample size (ESS>=400) and potential scale reduction factor (PSRF<1.1) of each model parameter.

To account for phylogenetic uncertainty, we ran MCMCglmm on the posterior sample of 100 trees, which is sufficient to obtain robust average probabilities ((Nakagawa & De Villemereuil, 2019); also see results and Fig. S1), and aggregated results of the nodes supported across trees. For each node, the estimated trait states are determined by comparing the liabilities and thresholds in each MCMC iteration. The aggregated iterations thus generate a posterior distribution of trait states which are used to estimate the probabilities of trait states at each node. The reconstructed posterior states of tip species were compared with their inputs to evaluate model performance.

### Phylogenetic PCA for ancestral reconstruction and shift inference

The threshold model estimates continuous liabilities and thresholds to model the evolution of ordered discrete traits. As a complementary analysis, we summarized diet divergence using phylogenetic principal component analysis (pPCA) and reconstructed ancestral positions in this dietary PCA space by modeling the first two principal components (PCs) as continuous variables in MCMCglmm. For easier PCA interpretation, we only used the three main diets (arthropods, nectar, fruits) in the pPCA. For each tree, we first conducted pPCA using the function *phyl.pca* in the R package phytools (Revell, 2024) with the lambda method (to account for the phylogenetic signal in the data), the correlation mode, and restricted maximum likelihood optimization. Then, the first two PCs were fitted to an MCMCglmm model as two continuous response variables following the Gaussian distribution and with non-fixed residual variance. The other model components were the same as described above. Each model was run with 10 independent chains and convergence was confirmed based on ESS and PSRF values. For this analysis, we only used the trees that showed the most-likely subfamily topology in (Rojas et al., 2016) and the genomic phylogeny. Results were aggregated across trees to estimate the mean posterior PC1 and PC2 scores which were projected to the input pPCA space to visualize the reconstructed locations of ancestral nodes.

In addition to node reconstructions, we also detected adaptive shifts of diets using the R package PhylogeneticEM (Bastide et al., 2018). For each of the trees having the most-likely subfamily topology, the first two phylogenetic PC scores estimated above were used as inputs and the scalar Ornstein–Uhlenbeck (OU) model was used to automatically detect adaptive shifts. We tested maximum 20 shifts with parallel alpha values, and the other parameters were kept as default. The automatically detected most likely number of shifts (according to the selection method LINselect) was used to extract branches that experienced a shift in the adaptive diet optimum.

## Results

### Input trees strongly support monophyly of bat (sub)families and genera

We first tested whether the phylogenetic trees, which were inferred from up to 31 gene sequences (Upham et al., 2019), support monophyly of each bat family. We found that all 10,000 trees support the monophyly of 20 among 21 bat families; the only exception is Mormoopidae which is paraphyletic in 98.95% of trees (although the two genera of this family, *Mormoops* and *Pteronotus*, are each monophyletic across all trees). We next tested the monophyly of the Phyllostomidae subfamilies and genera. All but one of the 10,000 trees supported the monophyly of the 11 subfamilies, with only a single tree indicating paraphyly of Desmodontinae regarding the position of Macrotinae. Among the 55 included phyllostomid genera, 49 are monophyletic across all 10,000 trees and four genera (*Choeroniscus*, *Tonatia*, *Vampyressa*, *Glyphonycteris*) are monophyletic in more than 99.7% trees. The other two genera, *Lonchophylla* and *Phyllostomus*, are paraphyletic in all trees, possibly due to the lack of data or species misidentification. In addition, all 10,000 trees supported the monophyly of the internal nodes of interest, including the MRCA of the superfamily Noctilionoidea (NM), the MRCA of Phyllostomidae and its most closely related family Mormoopidae (PM), the MRCA of all phyllostomids except for the basal subfamily Macrotinae (EB_Mic), the phyllostomid “early burst” node, and the MRCA of the four predominant fruit-feeding phyllostomid subfamilies (Stenodermatinae, Rhinophyllinae, Glyphonycterinae, Carolliinae; SRCG). In summary, the 10,000 input trees reliably captured the monophyly and major branching patterns of most bat families, Phyllostomidae subfamilies and genera.

When only considering the divergence of phyllostomid subfamilies, a total of 41 subfamily topologies were found in the 10,000 trees, all showing Desmodontinae (vampire bats) diverging before Micronycterinae (Fig. 1B). This position of Desmodontinae was only observed once with a combined dataset of molecular and morphological characters (Dávalos et al., 2014) and differs from all other molecular phylogenies (Fig. 1). Hence, this position is presumably an artifact due to the sparse data in the 31 gene matrix used to reconstruct these phylogenetic trees. Because the single origin of blood feeding in Desmodontinae is uncontested, with this being a binary trait only present in this subfamily, we excluded the three vampire bats from the following analyses to avoid incorporating phylogenetic bias in our reconstruction, leaving 621 bats in total and 176 bats in the Phyllostomidae-focused analyses. Nevertheless, including Desmodontinae and blood feeding does not affect our conclusions, as shown in Fig. S2 and S3.

To account for phylogenetic uncertainty in our analyses, we randomly sampled 100 trees for the following reconstructions. These 100 trees covered 11 of the 41 subfamily topologies, including the 6 most prevalent topologies with similar frequencies and represent phylogenetic conflicts in the literature (Fig. 1). Accordingly, except for cases with known phylogenetic conflicts, overall the 10,000 trees are in line with the current knowledge about the bat family phylogeny, and the 100 sampled trees effectively represented the phylogenetic uncertainty in these trees. The average probabilities plateaued after using more than 20 trees (Fig. S1), therefore the 100 trees were sufficient for robust estimates that account for phylogenetic uncertainties (using ∼50 trees has been suggested sufficient, (Nakagawa & De Villemereuil, 2019)).

### Diet reconstruction of Phyllostomidae

We first used the raw coding that coded species as complementary fruit-feeders even in case of sparse or ambiguous descriptions or when only trace amounts of fruits were detected in their diets. When using the raw coding, the Phyllostomidae-focused reconstructions showed high probabilities of complementary fruit feeding in the early burst node (94.1%) and the MRCA node of Phyllostomidae (88.1%; Fig. S4), providing support for ancestral omnivory. Complementary fruit feeding was even indicated in the MRCA node of Phyllostomidae and its most closely related arthropod-feeding family Mormoopidae (PM) with 18.3% probabilities, while outgroup families were reconstructed as strict arthropod feeding (Fig. S4). Since the high probabilities of ancestral complementary fruit feeding might be due to the highly inclusive coding of complementary fruit feeding in basal phyllostomid lineages, we tested how robust these results are when using the manually edited basal-insect coding, where the questionable cases were considered as absent fruit-feeding. Even when using the basal-insect coding, the MRCA of Phyllostomidae was still reconstructed with 44.4% complementary fruit feeding, and the early burst node was reconstructed with very high probabilities of complementary fruit (96.4%) and nectar (80.1%) feeding (Fig. 2). Since the manually edited basal-insect coding makes this analysis conservative, it provides strong support for the hypothesis of ancestral omnivory in Phyllostomidae. The two diet codings generated highly consistent results (Fig 2, Fig. S3) with major differences restricted to arthropod and fruit feeding at basal nodes of the family Phyllostomidae (Table 1). As expected, the subfamily Macrotinae was changed to strict arthropod feeding when using the basal-insect coding, reflecting changes in the input diets. The subfamily Lonchorhininae was not reconstructed with high probabilities of strict arthropod feeding probably because it is embedded among fruit- and nectar-feeding lineages in most trees (Fig. 1B), but its probability of strict arthropod feeding increased from 0.9% to 12.9% and the probability of complementary fruit feeding decreased from 89.0% to 57.2% when changing from raw to basal-insect coding (Table 1). Reconstructions of the subfamily Micronycterinae remained similar despite changes in diet codings, which accurately reflected the dietary diversity of this subfamily (Table S2). Hereafter we focused on the results from the basal-insect diet coding to avoid potential over-estimation of early-evolved complementary fruit feeding in Phyllostomidae.

**Figure 2.**
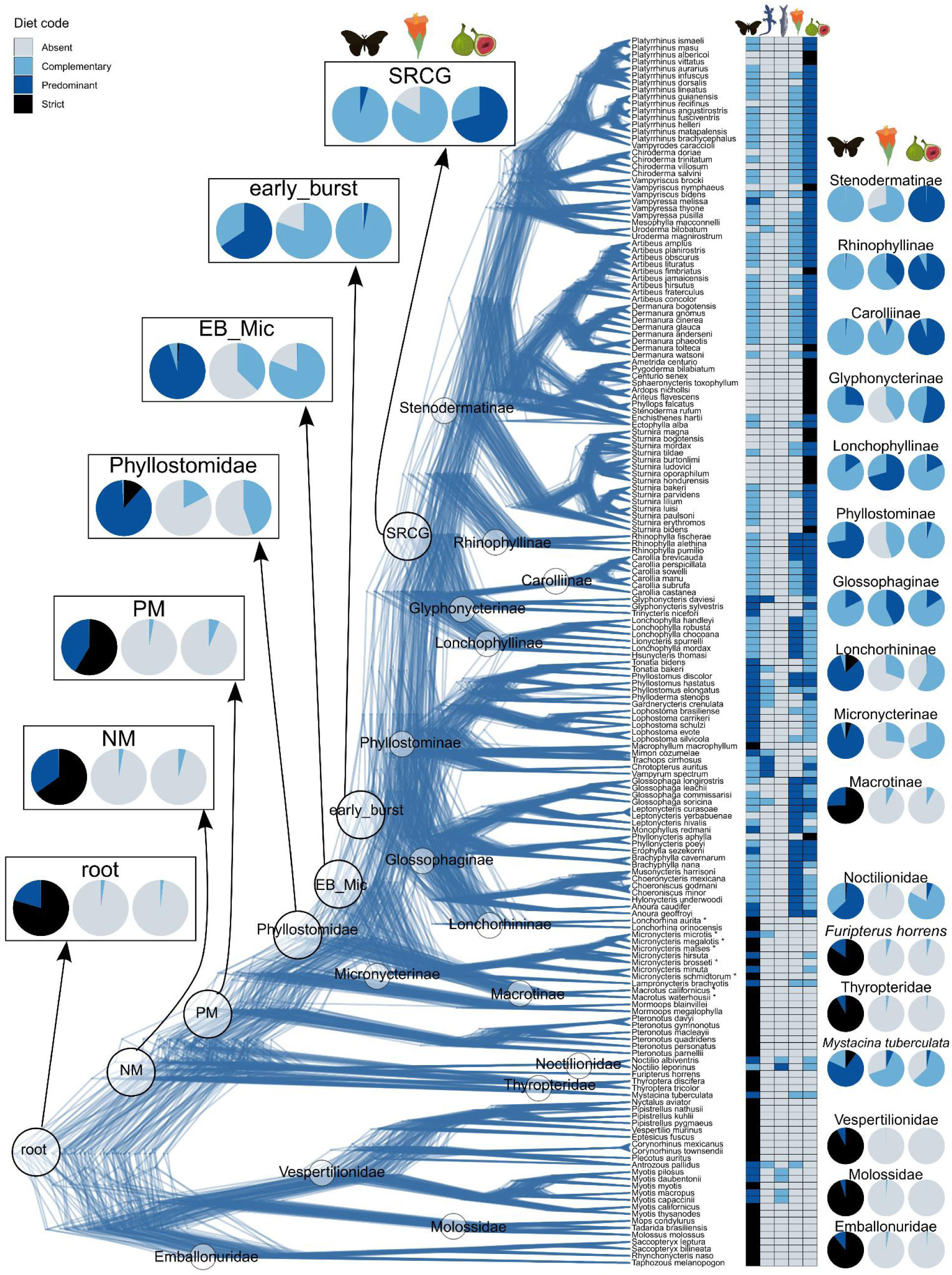
Diet reconstruction indicates early evolution of fruit feeding in Phyllostomidae. The densitree shows phylogenetic uncertainty captured by the 100 sampled trees of 176 bats. The heatmap shows the input states of five diets (arthropods, terrestrial vertebrates, fish, pollen and nectar, fruits) with the basal-insect coding. The manually re-coded tip species are labeled with asterisks. Pie charts show the reconstructed probabilities of internal nodes of interest (left) and the MRCA nodes of phyllostomid subfamilies and outgroup families (right). Single-species families (Mystacinidae, Furipteridae) are represented by posterior reconstructions of their tip species (*Mystacina tuberculata*, *Furipterus horrens*). The family Mormoopidae is not monophyletic in most trees; thus we omitted reconstructing its MRCA node. Node reconstructions of all five diets are given in Fig. S5. Fig. S3 shows a high similarity in the reconstructions when vampire bats are included and using the basal-insect coding.

**Table 1.**
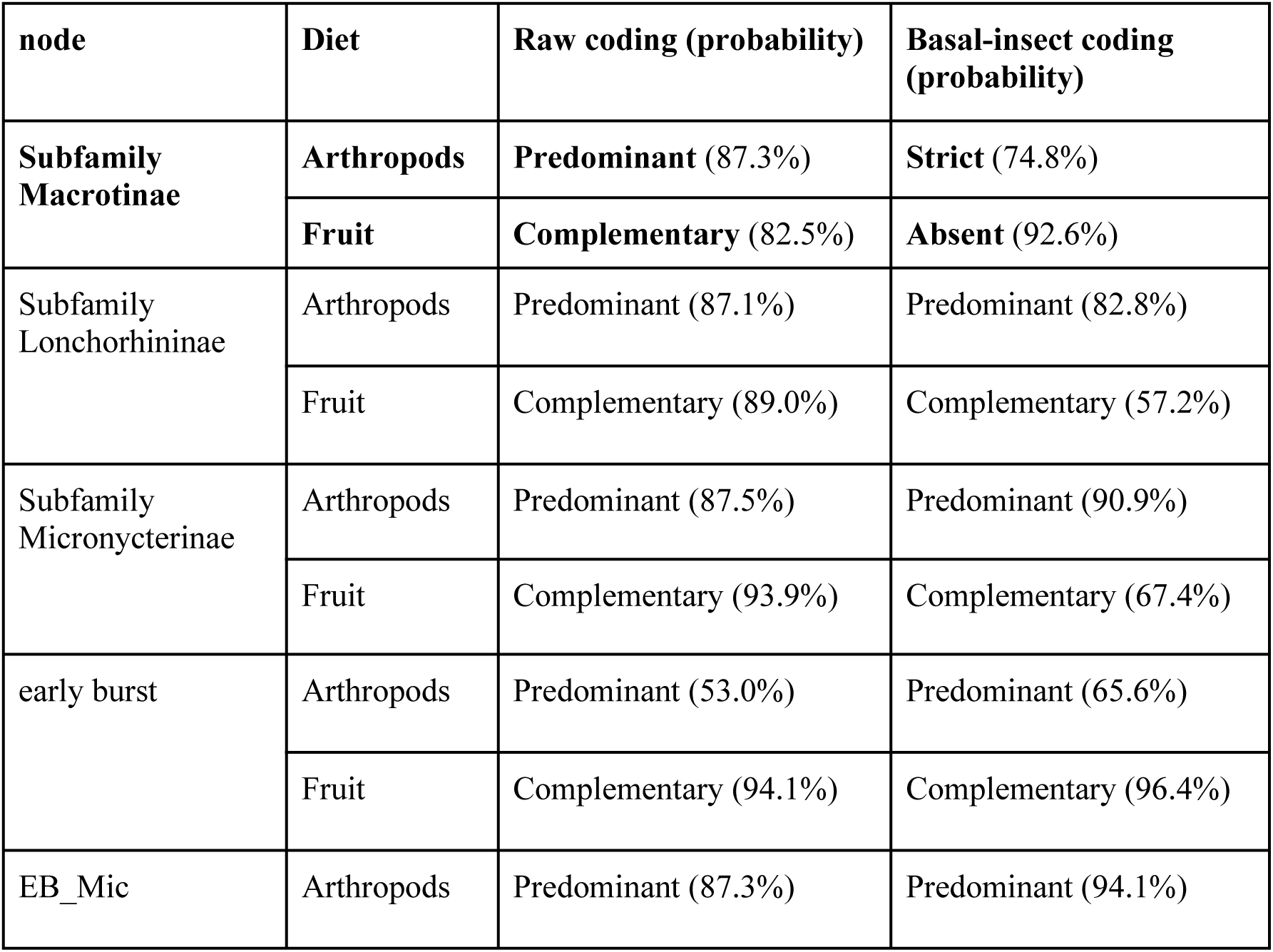
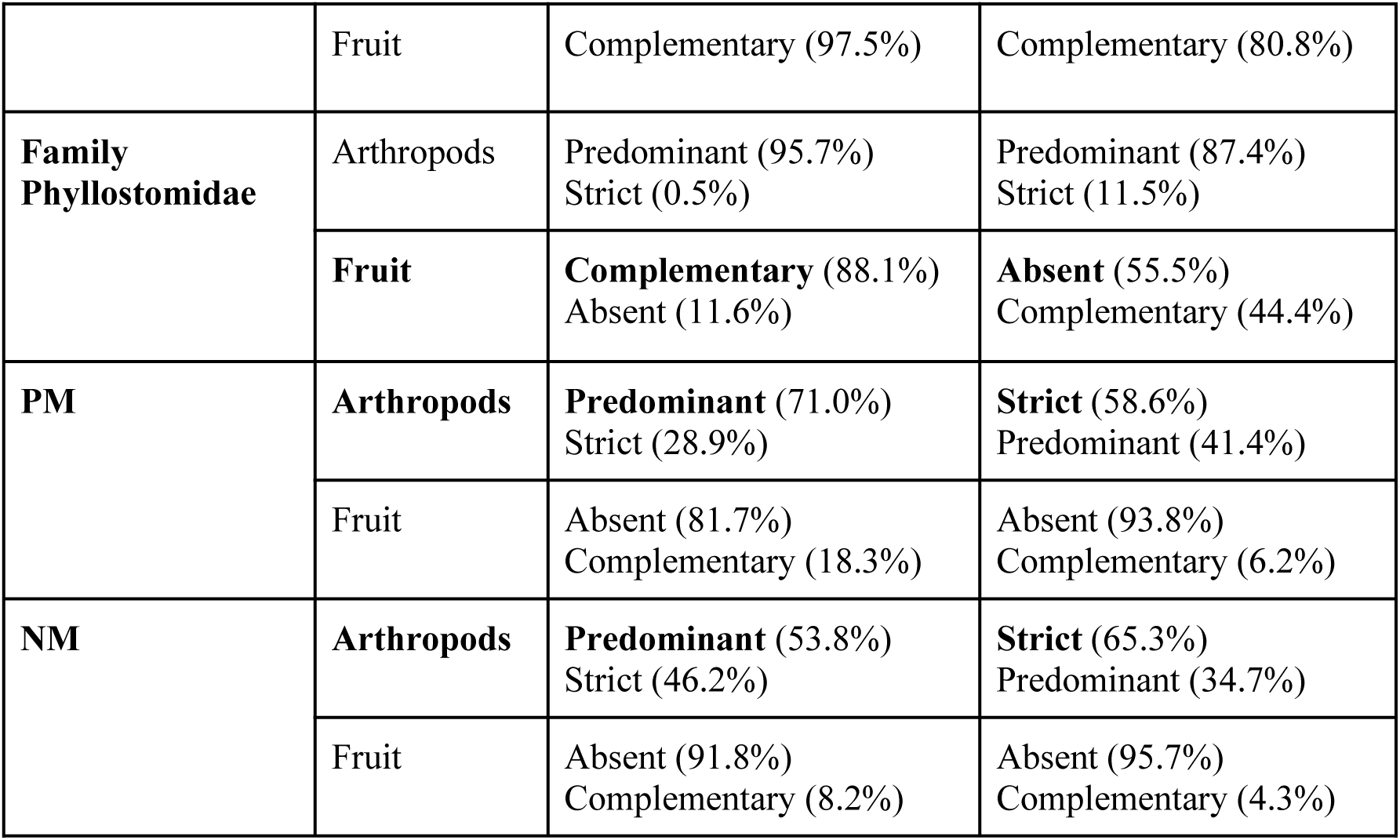
Node reconstructions using 176 bats and raw or basal-insect diet codings. The nodes and diets in bold had different reconstructions of the most likely states. Node labels are shown in Fig. 2: NM: the MRCA of the superfamily Noctilionoidea. PM: the MRCA of Phyllostomidae and Mormoopidae. EB_Mic: early burst plus Micronycterinae, i.e. the MRCA of all phyllostomids except for Macrotinae. Early burst: the MRCA of 8 subfamilies excluding Desmodontinae, Micronycterinae, and Macrotinae.

| node | Diet | Raw coding (probability) | Basal-insect coding (probability) |
| --- | --- | --- | --- |
| **Subfamily Macrotinae** | **Arthropods** | **Predominant (87.3%)** | **Strict (74.8%)** |
|  | **Fruit** | **Complementary (82.5%)** | **Absent (92.6%)** |
| Subfamily Lonchorhininae | Arthropods | Predominant (87.1%) | Predominant (82.8%) |
|  | Fruit | Complementary (89.0%) | Complementary (57.2%) |
| Subfamily Micronycterinae | Arthropods | Predominant (87.5%) | Predominant (90.9%) |
|  | Fruit | Complementary (93.9%) | Complementary (67.4%) |
| early burst | Arthropods | Predominant (53.0%) | Predominant (65.6%) |
|  | Fruit | Complementary (94.1%) | Complementary (96.4%) |
| EB_Mic | Arthropods | Predominant (87.3%) | Predominant (94.1%) |
|  | Fruit | Complementary (97.5%) | Complementary (80.8%) |
| **Family**<br>**Phyllostomidae** | Arthropods | Predominant (95.7%)<br>Strict (0.5%) | Predominant (87.4%)<br>Strict (11.5%) |
|  | **Fruit** | **Complementary** (88.1%)<br>Absent (11.6%) | **Absent** (55.5%)<br>Complementary (44.4%) |
| **PM** | **Arthropods** | **Predominant** (71.0%)<br>Strict (28.9%) | **Strict** (58.6%)<br>Predominant (41.4%) |
|  | Fruit | Absent (81.7%)<br>Complementary (18.3%) | Absent (93.8%)<br>Complementary (6.2%) |
| **NM** | **Arthropods** | **Predominant** (53.8%)<br>Strict (46.2%) | **Strict** (65.3%)<br>Predominant (34.7%) |
|  | Fruit | Absent (91.8%)<br>Complementary (8.2%) | Absent (95.7%)<br>Complementary (4.3%) |

To evaluate the performance of MCMCglmm models, we compared the inferred posterior probabilities of the diet variables of extant species to their input states. For the food items that most Phyllostomidae species feed on (arthropods, nectar, fruits), only 36 species had different input and posterior states and the differences are adjacent in the state order (Fig. S6), which is expected under the threshold model as liabilities close to the threshold would have similar chances of falling in either state divided by that threshold. Among all diets, the only case of non-adjacent states in the input and output was vertebrate feeding in *Glyphonycteris daviesi*: input predominant but output 45.6% probability for absent. Interestingly, there is no clear support for vertebrate feeding in the literature for this species, calling for further studies to clarify whether *Glyphonycteris daviesi* indeed feeds on vertebrates. Therefore, these results indicate overall consistent estimates and good performance of the MCMCglmm models.

### Reconstruction using the subfamily topology in the genome phylogeny

To test the effects of different topologies on the reconstructions, we estimated posterior probabilities using the 100 sampled trees but analyzed the 11 subfamily topologies separately. Reconstructions of internal nodes were overall consistent across subfamily topologies (Fig. S7). In particular, the early burst node was consistently reconstructed with high probabilities of complementary fruit (>90%) and nectar (>50%) feeding, and the MRCA of Phyllostomidae was consistently reconstructed with high probabilities of predominant arthropod feeding (>80%) and near 50% probabilities of complementary fruit feeding. However, some nodes showed higher variation in reconstructions across topologies, such as fruit feeding of the subfamily Lonchorhininae (Fig. S7) whose phylogenetic position varies in the 11 topologies (Fig. 1B).

Therefore, to further test the robustness of our diet reconstructions, we also estimated ancestral diets only using the trees that show the same subfamily topology in (Rojas et al., 2016) which is consistent with phylogenomic analyses. To this end, we considered all 23 trees out of the 10,000 trees that have this subfamily topology (only one of the 23 was included in the 100 sampled trees). Results aggregated across these 23 trees had the same most likely diet states with almost identical probabilities as the results from the 100 randomly sampled trees (Fig. 3A). As expected, minor changes were found in the MRCA of the subfamily Lonchorhininae which had relatively lower probabilities of strict arthropod feeding (7%; 13% in 100 trees) and higher probabilities of complementary fruit feeding (66%; 57% in 100 trees), consistent with its more recent divergence in the topology of the 23 trees (Fig. 3A).

**Figure 3.**
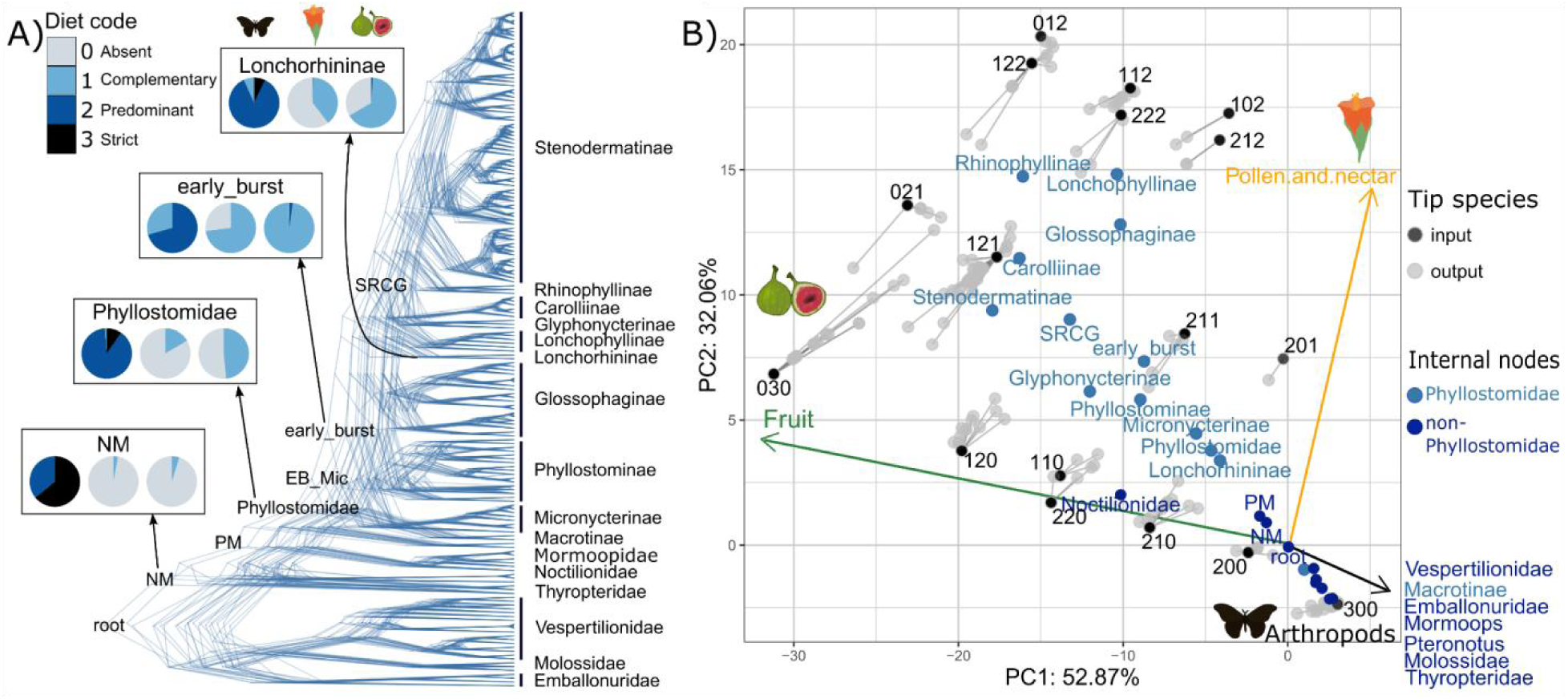
Diet reconstruction using 23 trees having the subfamily topology of the genome phylogeny. **A)** Reconstructions of discrete diet states using the threshold model. The densitree shows phylogenetic uncertainty in the 23 trees of 176 bats. Phyllostomid subfamilies and non-phyllostomid families are labeled on the right. Pie charts show the reconstructions of internal nodes of interest. **B)** Phylogenetic principal component analysis (pPCA) of the three main diets with projections of posterior PC reconstructions. The loadings and variance explained by the first two PCs are mean values of pPCA across 23 trees. Input pPCA scores of extant species are labeled with their diet states (in the order arthropods, fruit, nectar; basal-insect coding) and connected with their mean posterior pPCA scores. The internal nodes are projected using the mean reconstructed pPCA scores.

In addition to discrete diet states, we also inferred ancestral diets by reconstructing pPCA scores as continuous traits. Because pPCA plots are highly similar across trees, for easier visualization, we plotted the pPCA space using mean PC scores and mean diet loadings across trees, and projected the reconstructed posterior PC scores using the mean value estimated across MCMC interactions of the 23 trees. The reconstructed posterior PC scores of extant species were overall close to their corresponding inputs (Fig. 3B), except for several strict fruit-feeding species that were moved towards predominant fruit feeding and complementary nectar feeding in the posterior reconstructions, similar to the reconstructions of discrete states in extant species (Fig. S6). This is probably because transitioning to strict fruit feeding is a relatively rare event and the underlying liabilities were not estimated as extreme as the threshold given the input information in these models. On the other hand, strict arthropod-feeding species were all clustered in the pPCA space of athropod feeding, together with their ancestral nodes. Therefore, it is unlikely that a strictly insectivorous internal node would be reconstructed as omnivorous. Projections of internal nodes in the pPCA space indicated similar diets as the discrete-trait reconstructions (Fig. 3). For example, the early burst node was placed within the space of fruit and nectar feeding and close to the input state of predominant arthropod feeding and complementary fruit and nectar feeding (text label 211 in Fig. 3B), similar to the reconstructions of discrete diet states under the threshold model (Fig. 3A). The MRCA nodes of family Phyllostomidae, subfamily Micronycterinae, and subfamily Lonchorhininae were close to each other and closer to the origin but still in the space of fruit and nectar feeding (Fig. 3B), thus supporting an early evolution of these diets and ancestral omnivory.

To automatically and statistically detect adaptive diet shifts along the phyllostomid phylogeny, we ran PhylogeneticEM on the pPCA scores and the 23 phyllostomid phylogenetic trees that show the most likely subfamily topology. When using the PC scores estimated with the basal-insect coding, phylogeneticEM detected a median of 9 (range 6-16) shifts per tree. All of the 23 trees supported diet shifts on the branches leading to the early burst node and the subfamily Lonchorhininae, and more than 85% of trees supported diet shifts on the branches leading to the subfamily Glyphonycterinae, the MRCA of four predominant fruit-eating subfamilies (SRCG), and the family Noctilionidae (Fig. S8A). When using the PC scores estimated with the raw coding, phylogeneticEM detected median 7 (range 3-17) shifts per tree. In 87% (20 of 23) of trees, diet shifts were detected on the branches leading to the family Phyllostomidae, the subfamily Lonchorhininae, and the subfamily Glyphonycterinae. Diet shifts were also detected on the branch leading to the subfamily Stenodermatinae in 65% trees, on the branch leading to the family Noctilionidae in 48% of trees, and on the branch leading to SRCG in 35% of trees, but the branch leading to the early burst node was only inferred to have diet shifts in 5% of trees (Fig. S8B). Accordingly, the earliest shift of adaptive diet optimum in phyllostomid bats was indicated in the common ancestor of Phyllostomidae when using the raw coding and in the early burst stage when using the basal-insect coding.

### Diet reconstruction of all bats

Lastly, we expanded our analyses to all bat families, covering a total of 621 bat species (excluding the three vampire bats). Using the same 100 sampled trees and the basal-insect coding, we obtained highly similar reconstructions for Phyllostomidae and the other families in the superfamily Noctilionoidea (Fig. 4). In addition to Phyllostomidae, another three families were also reconstructed with ancestral fruit feeding: Noctilionidae, Mystacinidae, and Pteropodidae (Old World fruit bats). Similarly, nectar feeding was reconstructed in the families Phyllostomidae, Mystacinidae, and Pteropodidae. In addition, families Noctilionidae and Megadermatidae were reconstructed with ancestral complementary fish feeding, and Nycteridae and Megadermatidae were reconstructed with ancestral feeding on terrestrial vertebrates. Despite the indicated ancestral diet diversity in these families, ancestral insectivory was still strongly supported for all bats (order Chiroptera) and the two suborders (Yinpterochiroptera and Yangochiroptera).

**Figure 4.**
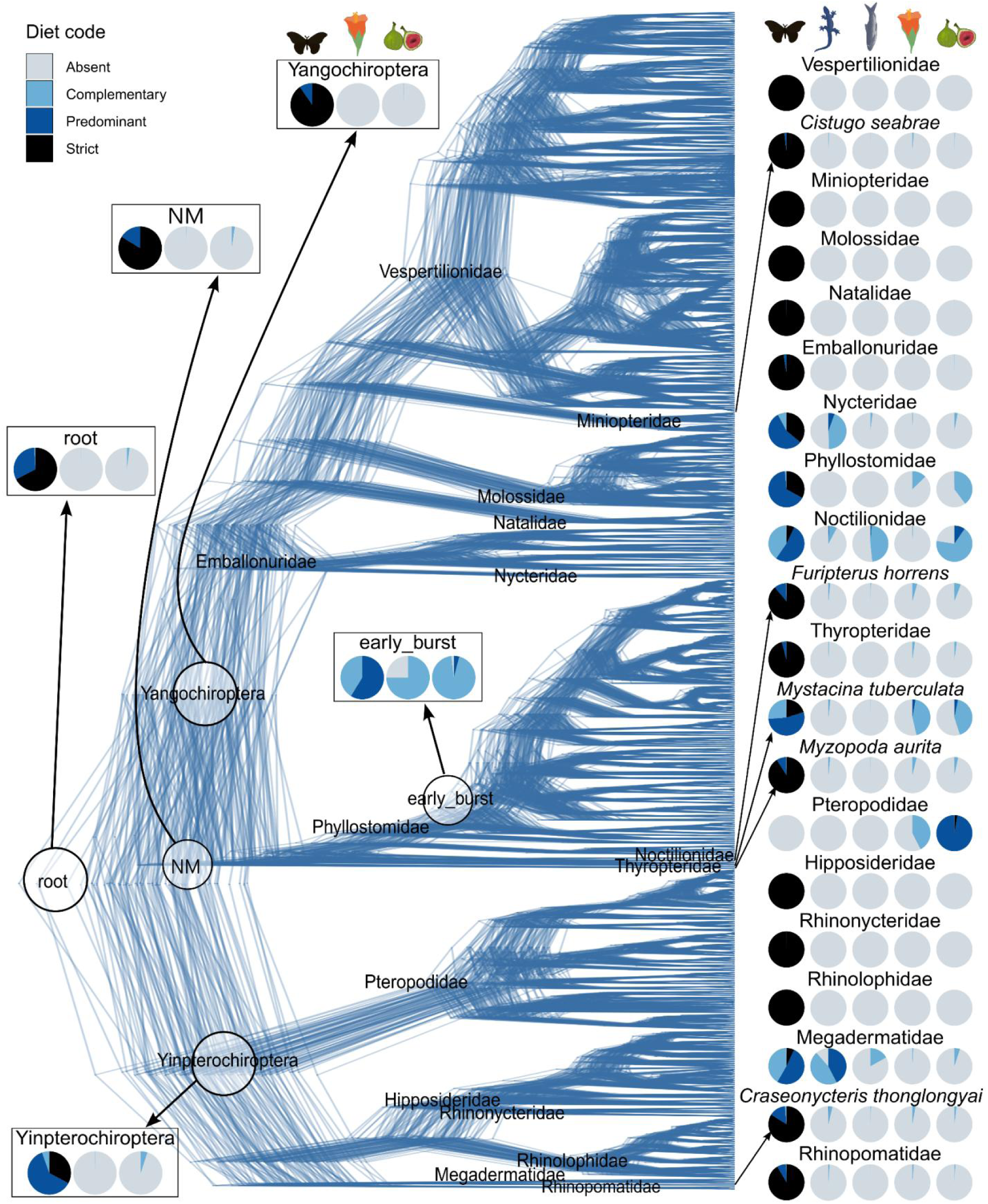
Diet reconstruction of all bat families covering 621 species. The densitree shows phylogenetic uncertainty captured by the 100 sampled trees of 621 bats. Pie charts on the right show the reconstructions of MRCA nodes of 21 bat families. Reconstructions of the five single-species families (Cistugidae, Furipteridae, Mystacinidae, Myzopodidae, Craseonycteridae) are the posterior probabilities of their extant species. Pie charts on the left show the reconstructions of the internal nodes of interest. NM refers to the MRCA node of the superfamily Noctilionoidea (same in the above trees) and does not include the family Myzopodidae which was grouped with Noctilionoidea when constructing these mammalian trees (Upham et al., 2019) but this fixed position contradicts the other bat phylogenies (Teeling et al., 2018).

## Discussion

Accurate inferences of ancestral diets in the ecologically diverse family Phyllostomidae provide a key foundation for understanding their adaptive radiation and testing the hypothesis of ancestral generalism. In contrast to previous work (Grossnickle et al., 2024; Rojas et al., 2011), our diet reconstructions consistently indicate early evolution of complementary fruit feeding in Phyllostomidae, which supports the hypothesis of ancestral generalism. The common ancestor of Phyllostomidae was most likely predominant (rather than strict) arthropod-feeding, with fruits being the most likely alternative complementary diet. This conclusion was robustly drawn regardless of the input diet coding, underlying phylogenetic topologies, or the inclusion of vampire bats. Even when using the basal-insect coding, the probabilities of complementary fruit feeding were close to 50% at the MRCA node of Phyllostomidae, indicating that the underlying liabilities were close to the threshold between absent and complementary states, which is expected when a trait first evolved. Whereas complementary fruit feeding probably did not evolve before the divergence of Phyllostomidae (<10% probability at the PM node using the basal-insect coding), it likely fully evolved at the early burst node (>90% probability). In the more rigorous analyses of adaptive shifts, the basal-insect coding detected an adaptive shift on the branch leading to the early burst node but not the branch leading to the family Phyllostomidae, indicating that the complementary fruit feeding that evolved before the early burst (e.g., in the subfamily Micronycterinae and the MRCA of Phyllostomidae) was an expansion of the dietary niche but was not enough to represent a different adaptive optimum. Therefore, we suggest that complementary fruit feeding has at least started to evolve in the ancestral lineage of Phyllostomidae, possibly as a mechanism of opportunistic omnivory to cope with food fluctuation, similar to some extant species that are also generalists (Rex et al., 2010). Our results support the hypothesis of ancestral omnivory and generalism, which may have helped to expand ancestral ecological niches early in the phyllostomid adaptive radiation (Davies et al., 2020; Freeman, 2000; Hall et al., 2021; Potter et al., 2021).

High probabilities of complementary fruit-feeding and nectar-feeding at the early burst node support a previous study by (Grossnickle et al., 2024) that used molar traits to infer that phyllostomid bats have invaded these alternative feeding habits at the early burst stage. Our pPCA pattern of the three main diets is also highly consistent with the PCA of diet-related molar traits (Grossnickle et al., 2024). However, the previous diet reconstruction only indicated nectar feeding at the early burst node (Grossnickle et al., 2024), whereas our results showed higher probabilities of complementary fruit feeding than nectar feeding in ancestral nodes, indicating that fruit feeding evolved before nectar feeding. This is expected because nectarivory is more derived and specialized than frugivory. In other words, the adaptive valley between insectivory and frugivory is probably easier to cross than the adaptive valley between insectivory and nectarivory, making fruit feeding more likely to be the first ancestral niche expansion. However, by the early burst stage, both complementary fruit and nectar feeding probably have fully evolved, and this omnivory has been maintained in many subsequently diverged subfamilies despite increased specialization in fruit and nectar feeding. For example, the oldest plant-visiting bat fossil (in the subfamily Lonchophyllinae) was indicated to be omnivorous feeding on insects, fruits, and nectar (Dávalos et al., 2014; Yohe et al., 2015), consistent with our reconstructions. The subsequent diversification within fruit- and nectar-feeding lineages represents either specialisation (e.g., in the fruit-eating subfamily Stenodermatinae), or niche partitioning while maintaining some level of generalism (e.g., in the subfamily Glossophaginae).

Reconstructions of all bat families indicated that fruit feeding evolved at least four times in the MRCA nodes of the families Phyllostomidae (227 extant species in central and south America), Pteropodidae (202 species in the Old World), Mystacinidae (one species endemic in New Zealand), and Noctilionidae (two species in central and south America). A few species in the families Emballonuridae and Vespertilionidae were inclusively coded as complementary fruit or nectar feeding but the MRCA nodes of these families were strict insect feeding. The single species in Mystacinidae, *Mystacina tuberculata*, predominantly feeds on arthropods but has been found to also eat fruits and nectar (Arkins et al., 1999; Carter & Riskin, 2006; Daniel, 1976; McCartney et al., 2007). Similar omnivorous diets have also been suggested for fossil species of Mystacinidae which showed a decline in morphological diversity since the Miocene (Hand et al., 2015, 2018). Both species in Noctilionidae were inclusively coded as complementary fruit feeding (Rojas et al., 2018), although occasional fruit feeding was mostly found in *Noctilio albiventis* (Aranguren et al., 2011; Gonçalves et al., 2007) and lacks solid literature support in *Noctilio leporinus*. Therefore, our reconstruction might overestimate the proportion of fruit feeding in the MRCA of Noctilionidae. However, even when using the inclusive raw coding, the MRCA of the superfamily Noctilionoidea (NM, including three ancestral fruit-feeding families Phyllostomidae, Noctilionidae, and Mystacinidae) was consistently reconstructed with very low probabilities of fruit feeding and high probabilities of strict arthropod feeding. Therefore, our results indicate that fruit feeding evolved repeatedly and independently in these families.

Although several families have evolved complementary fruit feeding, predominant and strict fruit feeding has been found only in the specious families Phyllostomidae and Pteropodidae. The other families have very low species diversity despite ancestral omnivory. Therefore, ancestral generalism, represented by ancestral complementary fruit and nectar feeding, may not be sufficient to promote speciation despite expanding ancestral dietary niches. Subsequent niche partitioning is also required for successful adaptive radiations.

In addition to fruit and nectar feeding, our reconstructions also supported multiple independent evolutionary origins of carnivory in bats. Feeding on terrestrial vertebrates was reconstructed in the ancestral nodes of families Nycteridae and Megadermatidae, and was found in several species in the families Pteropodidae, Vespertilionidae, and Phyllostomidae. Similarly, fish feeding was reconstructed in the ancestral nodes of families Noctilionidae and Megadermatidae and was found in several species in the genus *Myotis* (MRCA of *Myotis* was strict arthropod feeding; note that the predominant fishing bat *Myotis vivesi* was not included in the input trees). Some studies suggested that carnivory in bats is one extreme of animalivory (the other extreme being insectivory) and is mainly driven by larger body size (Giannini & Kalko, 2005; Santana & Cheung, 2016). Arthropod feeding and vertebrate feeding might represent overall similar feeding habits and ecological niches, which might explain why the evolution of vertebrate feeding did not promote speciation in the corresponding families.

Except for ancestral diet diversity reconstructed in the above families, most of the 21 bat families were found with strict arthropod-feeding in their MRCA nodes. Strict arthropod feeding was also supported in the common ancestor of all bats and the two suborders (Yinpterochiroptera, Yangochiroptera), which is consistent with the canonical hypothesis that bats have an insectivorous most recent common ancestor (Gunnell & Simmons, 2005; Simmons et al., 2008). A recent study suggested an omnivorous common ancestor of bats based on sugar responses of the reconstructed ancestral sweet taste receptors (Tas1r2 and Tas1r3; (Li et al., 2023)). However, responses of sweet taste receptors may not be sufficient to indicate diet preferences. For example, the same study did not find sugar responses of the sweet taste receptor in fruit-feeding phyllostomid bats (Li et al., 2023), and a study in songbirds found sugar responses across species with diverse diets as they retained an ancestral sugar-sensing ability (Toda et al., 2021). Accordingly, we suggest that bats most likely evolved from an insectivorous common ancestor and independently gained fruit feeding in at least four families.

## Conclusion

Our study supports the hypothesis of ancestral generalism (i.e., omnivory) in the bat family Phyllostomidae that displays an unparalleled dietary diversity in mammals. We showed that complementary fruit feeding has started to evolve in the common ancestor of Phyllostomidae, and both complementary fruit and nectar feeding have fully evolved by the early burst stage, as was also indicated by the diet-related molar traits. Different from previous diet reconstructions, our results show that complementary fruit feeding evolved earlier than nectar feeding. In addition, we provide a comprehensive picture of dietary evolution in all bat families and supported ancestral insectivory of the order Chiroptera and its two suborders. Independent evolution of ancestral fruit feeding was found in four families, but only the two families (Phyllostomidae and Pteropodidae) harboring predominant and strict fruit feeders show high species diversity. Therefore, ancestral generalism may be a precondition of but does not necessarily lead to adaptive radiation which also requires subsequent niche partitioning.

## Acknowledgment

This work was supported by the German Research Foundation (HI1423/6-1) and the LOEWE-Centre for Translational Biodiversity Genomics (TBG) funded by the Hessen State Ministry of Higher Education, Research and the Arts (LOEWE/1/10/519/03/03.001(0014)/52). We thank Dr. Leon Hilgers for the insightful comments on the manuscript.

## Author Contributions

Conceptualization: XY and MH. Data curation and analyses: XY. Funding and supervision: MH. Investigation: XY and MH. Methodology: XY, DGK, MH. Validation: XY, DGK, MH. Visualization: XY. Original draft: XY. Review and editing: XY, DGK, MH.

## Data and Code Availability

Input phylogenetic trees are available from VertLife (http://vertlife.org/phylosubsets). Index of the subset of trees used in our reconstructions and the R scripts used in this study are provided on GitHub (https://github.com/xuelingyi/Bat_Diet_Reconstruct).

