## Supplementary material for "Comprehensive phylogenetic reconstructions support ancestral omnivory in the ecologically diverse bat family Phyllostomidae": FigS1-S8, Table S1-S2

The supplement contains

- Figures S1 to S8; Figure S5 is provided as an independent PDF file (legend below).
- Tables S1 and S2 are provided as sheets in an Excel file.

**Table S1. The input bat diets.**
The column "vertlife_name" gives tip names in the input trees and the column "ScientificName" gives updated species names (used in plots). Six species had duplicated scientific names after the taxonomic update. Some individuals of the same species were not sisters in the phylogenetic trees, indicating misidentification, but they were always grouped in the same subfamily and thus should not impact our conclusions. The column "conservative_coding" labels the species re-coded in the basal-insect coding. The column "dataset" indicates whether the species was included in the Phyllostomidae-focused analyses (sif176) and/or the all bat analyses (bat621). The three vampire bats are also labeled.

**Table S2. The diet review of predominantly insectivorous phyllostomid lineages.**
Species that do not have sufficient support for fruit-feeding were re-coded as strict insectivorous in the basal-insect diet coding.

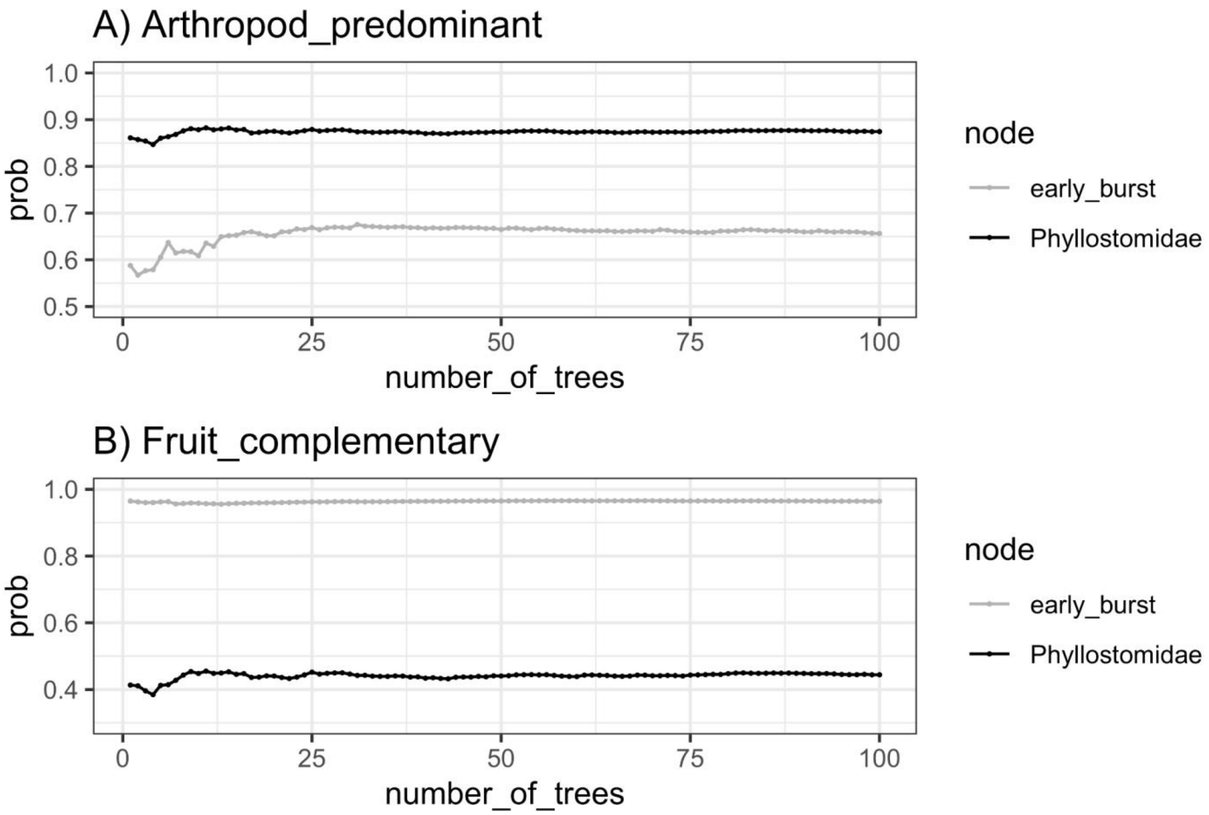


**Figure S1. Average probabilities of 1-100 of the sampled trees for the two focal nodes and diet states.**

The results are from analyses of 176 bats using the basal-insect diet coding. Each dot represents average probabilities using the corresponding number of trees.

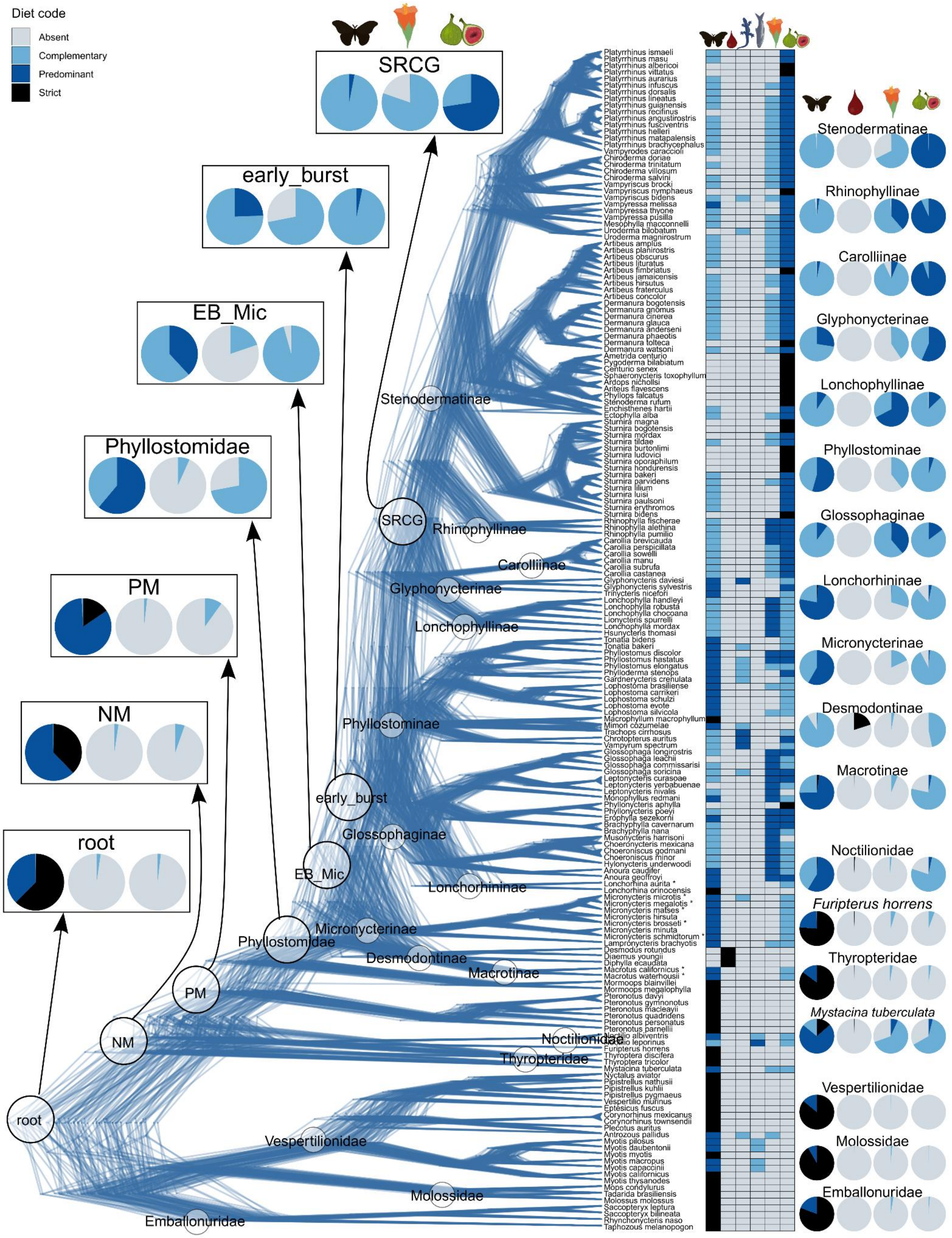


**Figure S2. Node reconstructions with the raw coding and including vampire bats**.

The densitree shows phylogenetic uncertainty captured by the 100 sampled trees of 179 bats. The heatmap shows the input states of six diets (arthropods, blood, terrestrial vertebrates, fish, pollen and nectar, fruits) with the raw coding. Pie charts show the reconstructions of internal nodes of interest (left) and the MRCA nodes of phyllostomid subfamilies and outgroup families (right). Single-species families (Mystacinidae, Furipteridae) are represented by posterior reconstructions of their tip species (*Mystacina tuberculata*, *Furipterus horrens*). The family Mormoopidae is not monophyletic in most trees and thus its MRCA node was not reconstructed. To improve the MCMCglmm fit for blood feeding, in addition to the three vampire species, their MRCA and internal nodes of the subfamily Desmodontinae were also set as strict blood-feeding in the input. The reconstructions showed 80% strict blood-feeding for the species *Desmodus rotundus* and *Diaemus youngii*, and 66% strict blood-feeding for the species Diphylla ecaudata, but only 20% strict blood feeding for the MRCA node of the subfamily Desmodontinae. This most likely reflects the limited power of MCMCglmm in modeling a binary trait that only evolved in a monophyletic clade of three species. Reconstructions of the main diets (arthropod, fruits, nectar) are highly consistent between results including and excluding vampire bats (Fig. 2, Fig. S2, S3, S4), with the only difference being the arthropod feeding in the early burst node (most likely complementary changed to predominant after excluding vampire bats) and in the PM node (most likely predominant changed to strict after excluding vampire bats). Complementary nectar and fruit feeding remained the most likely states at the early burst and phyllostomid MRCA nodes but the probabilities were lower when including vampire bats. In summary, reconstructions including vampire bats tend to indicate less of the main diets in ancestral nodes (Fig. S2, S3) because the early-diverged vampire bats have absent arthropod, nectar, and fruit feeding. However, the overall results and conclusions are consistent, and so in the main text we focus on the analyses excluding vampire bats.

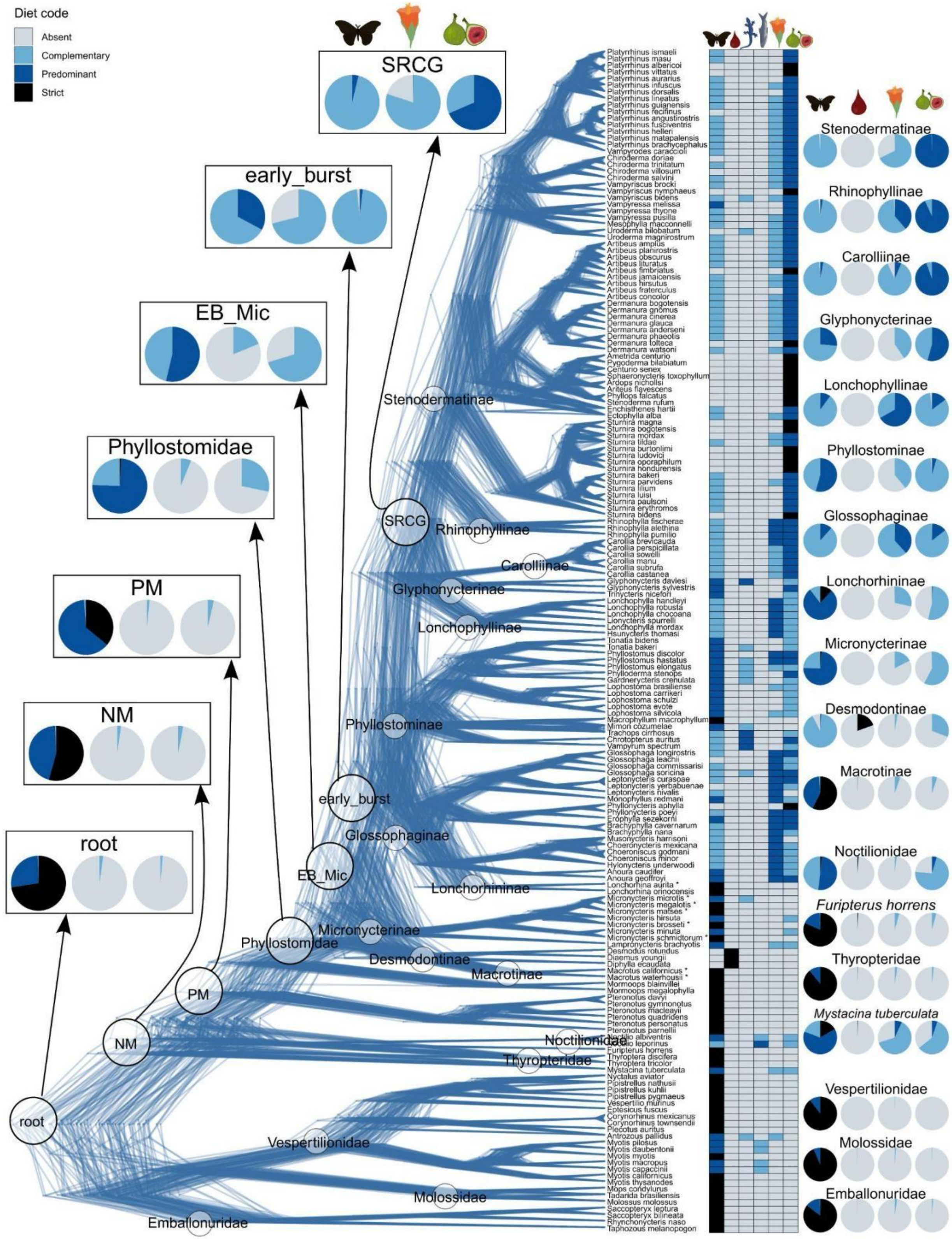


**Figure S3. Node reconstructions with the basal-insect coding and including vampire bats.** Tip species labeled with asterisks are manually re-coded. Same legend as in Fig. S2.

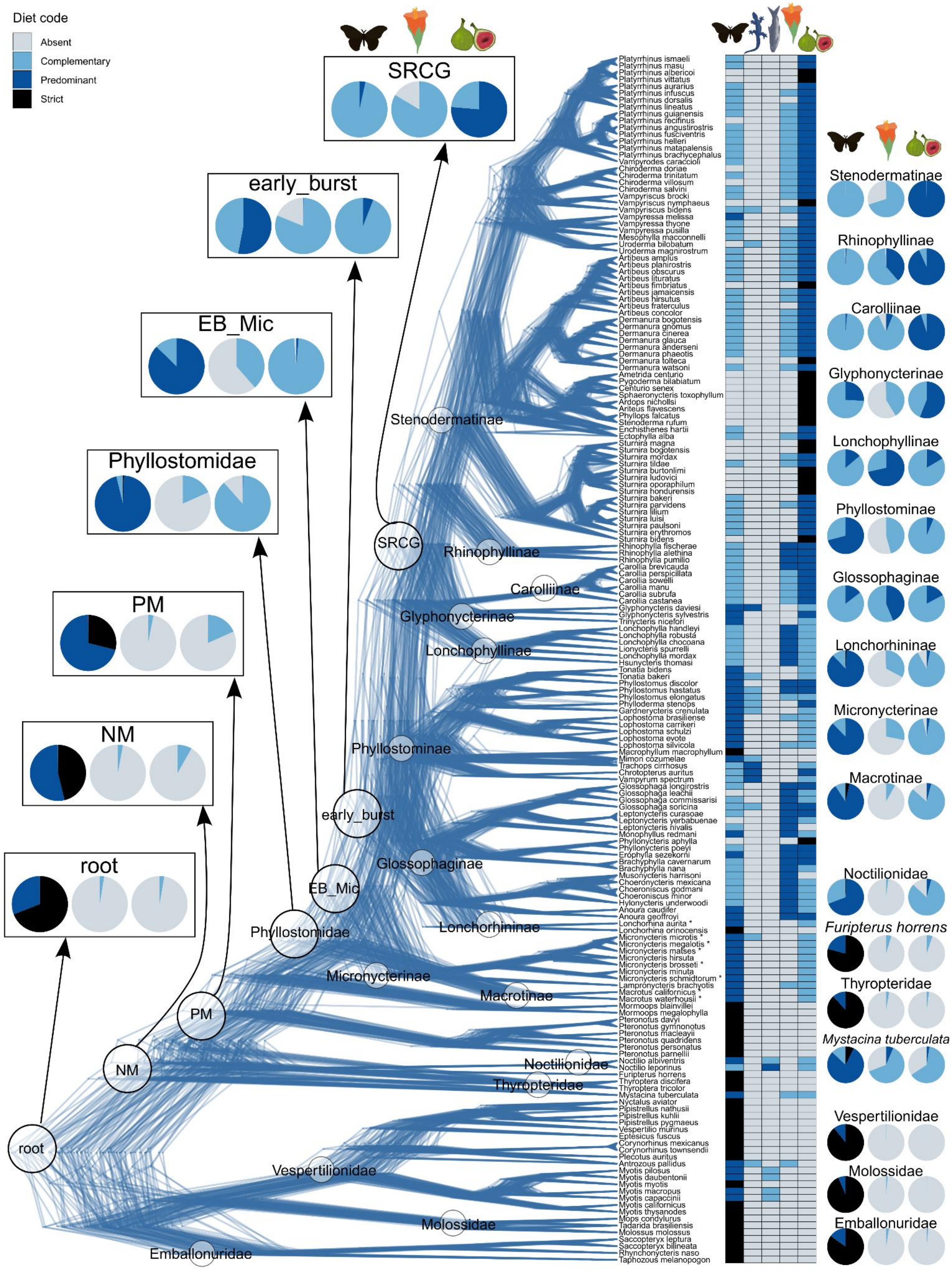


**Figure S4. Node reconstructions in 176 bats using the raw diet coding**.
Same legend in Fig. 2.

**Figure S5. Node reconstructions of all 5 food items in 176 bats using the raw and basal-insect diet codings**.

Each pie chart is labeled with its most likely state and the corresponding probability. [independent PDF]

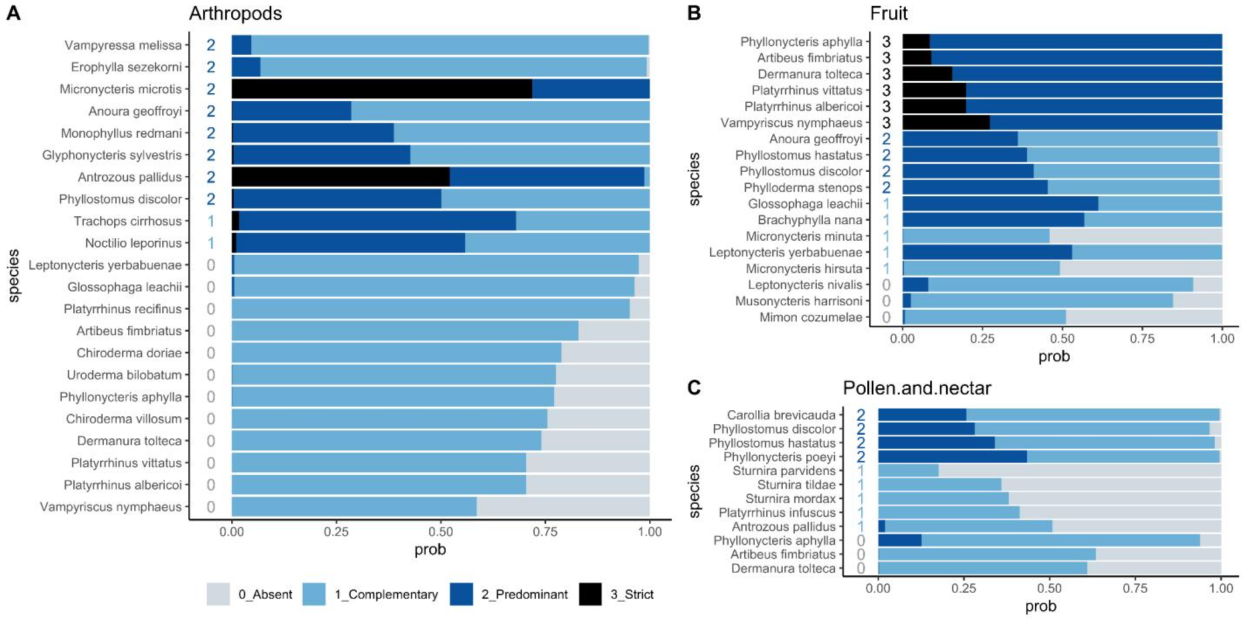


**Figure S6. Tip species whose most likely posterior states differ from the input states using the basal-insect coding.**

The results are from analyses of 176 bats using the basal-insect diet coding. Each horizontal bar corresponds to one species. Numbers on the left represent the input state and bar components represent posterior probabilities of each state.

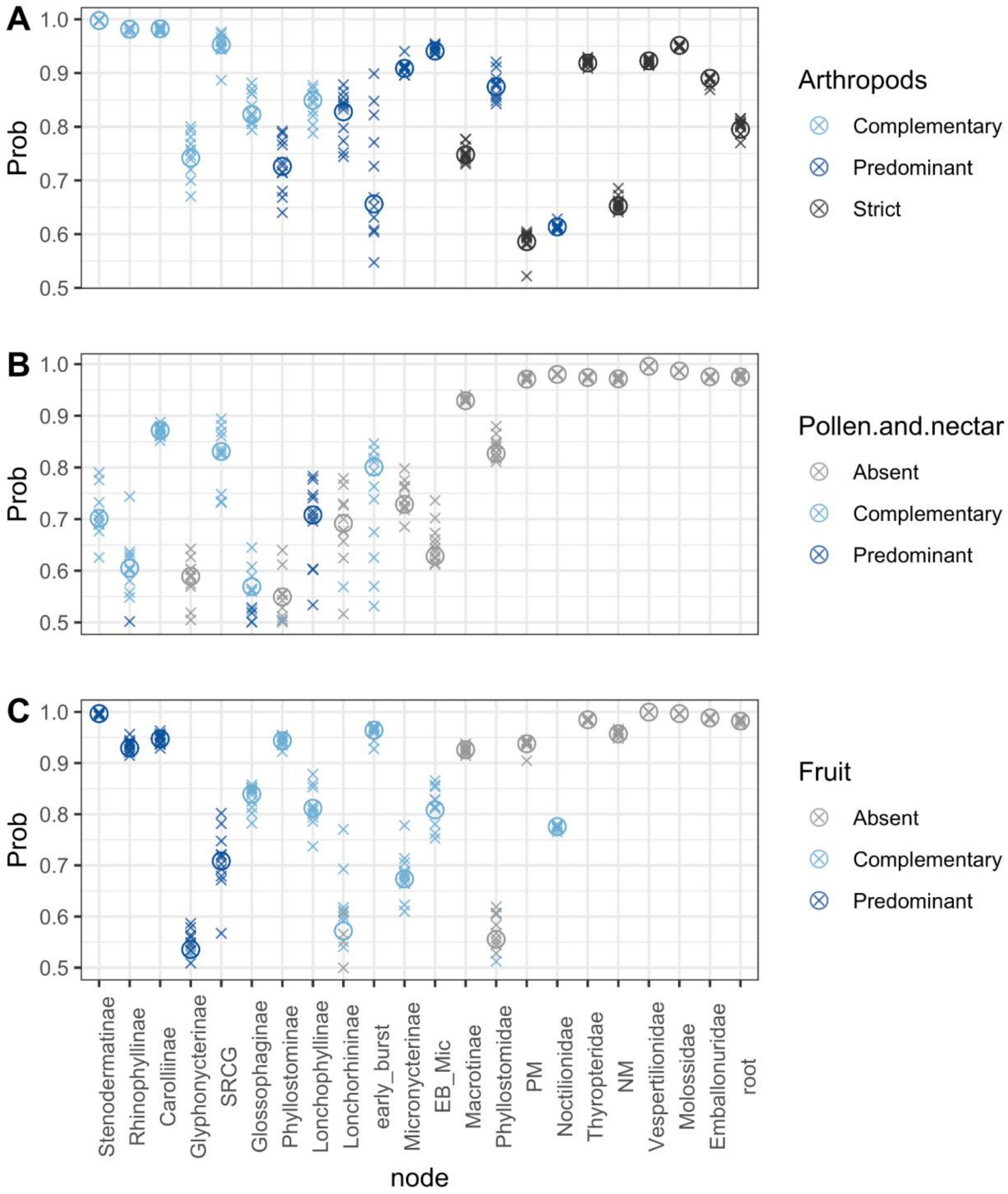


**Figure S7. Reconstructions of the most likely states in each subfamily topology.**
The results are from 176 bats using the basal-insect diet coding. Each cross represents the most likely posterior state based on the results aggregated across trees having the same subfamily topology (total 11 topologies in the 100 sampled trees). Circles represent the most likely posterior state estimated by aggregating all 100 trees.

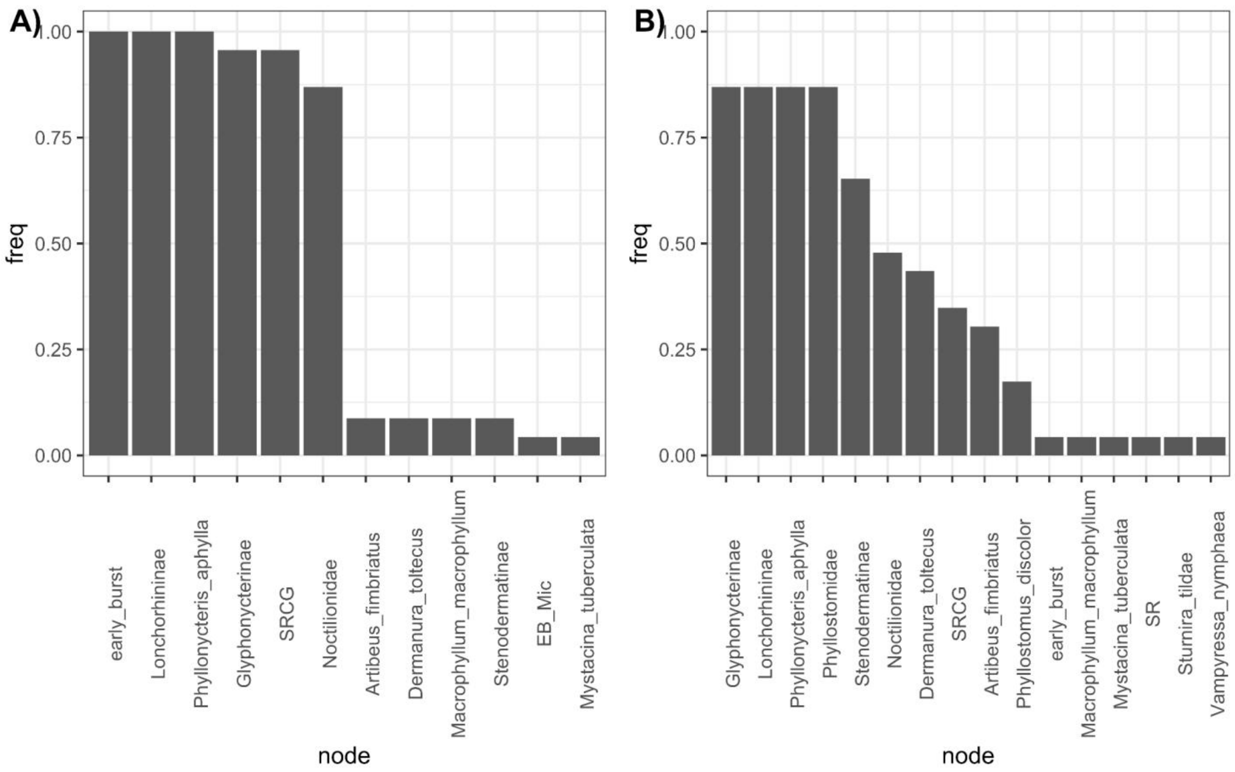


**Figure S8. The most frequently detected adaptive shifts leading to ancestral nodes of Phyllostomidae.**

The input pPCA scores were estimated using three main diets (arthropods, fruit, nectar) with **A)** the basal-insect coding and **B)** the raw coding. Only the labelled nodes are presented, whereas the non-labelled nodes all have low frequencies (<0.15 in the basal-insect coding, <0.10 in the raw coding) and may not be shared across trees.
